# Computational Recipe for Designing Antibodies against the Ebola Virus

**DOI:** 10.1101/2020.04.04.024430

**Authors:** Amir Barati Farimani, Narayana R. Aluru, Emad Tajkhorshid, Eric Jakobsson

## Abstract

A conceptual basis for antiviral therapy is to deliver a synthetic antibody that binds to a viral surface protein, and thus prevents the virus from deploying its cell-entry mechanism. The fast and untraceable virus mutations take lives of thousands of people before the immune system can produce the inhibitory antibody. In this paper, we devised a computational recipe to predict both the viral escape mutations and the possible inhibitory synthetic antibodies. We combined bioinformatics, structural biology, and molecular dynamics (MD) simulations to explore the most likely viral mutations and the candidate antibodies that can inhibit those escape mutations. Specifically, using the crystal structures of the Sudan and Zaire Ebola viral GPs in complex to their respective antibodies (ABs), we have performed an extensive set of MD simulations, both on the wild-type structures and on a large array of additional complexes designed and generated through combinatorial mutations. We discovered that our methods enabled the successful redesign of antibody sequences to essentially all likely glycoprotein mutations. Our findings and the computational methodology developed here for general antibody design can facilitate therapy of current and possibly next generations of viruses.

**Significance of the Manuscript:** This manuscript has high significance both methodologically and in potential biomedical application. In methodology, the manuscript combines molecular dynamics, Monte Carlo calculations, and bioinformatics in a novel way to simulate the evolutionary arms race between an evolving viral coat protein and a counter-evolving antibody against the virus. This simulation is shown to provide a method for designing a synthetic antibody against the newly emerging viral strains. This work is done in the context of ongoing work in other laboratories in which cells can be induced to produce synthetic antibodies and those synthetic antibodies can be edited (via, for example, CRISPR) to have an arbitrary sequence in the region that binds the viral coat protein. Putting those experimental methods together with the computational methods we present in this paper has the potential to provide a important approach to produce antibodies-on-demand against evolving viruses.

## Introduction

The ability to produce antibodies specific to predefined biomolecular targets was a landmark development in biological research and potential therapy.^1^ Using this ability as a foundation, therapeutic antibodies have been developed.^2^ It is possible to produce antibodies with a completely synthetic antigen-binding region, by inserting synthetic DNA into the appropriate space in the antibody-encoding gene(s).^3^ Nature and structural biologists have provided us with a proof of concept for such engineering, by providing us with definable sequences and structures for viral proteins, e.g., Ebola glycoproteins(GP), bound to antibodies that enabled the host to survive the disease.^4^ On the other hand, nature makes our task harder by enabling the virus to evolve in such a way as to neutralize the effect of the antibodies.^4^ An outline of how the evolutionary arms race proceeds between Ebola virus glycoprotein vs. antibodies from the host immune system is provided by sequences and structures for multiple glycoprotein-antibody complexes. However this description of the glycoprotein-antibody competition does not in itself lead to a predictive model for how the virus will evolve and what change in the antibody will be effective against the evolved viral protein. The purpose of this paper is to provide such a model.

Among the structurally resolved viral protein-antibody complexes, Ebola is an important target due to persistent recurring outbreaks and very high case fatality rate.^5^ A couple of promising computational efforts have been carried to virtually screen possible small molecule candidates to inhibit the Ebola VP40 viral protein.^6^ Miller et al. also mapped the Ebola virus escape mutations and created a watchlist.^7^ More detailed investigations, however, are needed on how to design an effective antibody before a new strain appears, in order to prevent another outbreak disaster. The purpose of the present paper is to use the tools of computational biology combined with the above-cited sequences and structures to construct a predictive model for how the Ebola virus is likely to evolve, and what alterations in the sequence of binding region of the antibody would most effectively counter the viral mutation(s) and restore the ability of the antibody to bind the glycoprotein. An effective approach to predict likely mutations is to use existing statistical data on the likelihood of particular substitutions. These data are typically embodied in a “substitution matrix” in which each element corresponds to a relative probability of an amino acid substitution.^8^ A reasonable tool to predict effective responses to viral mutations is to use existing statistical data on amino acids that interact favorably at the protein-protein interface between the viral protein and the antibody.^9^ Finally, molecular dynamics simulations of the mutated glycoprotein-antibody complex can be used to test the statistical prediction by computing the structural and energetic consequences of the postulated mutations.^10^

The body of this paper will describe the implementation of this computational strategy.

## Methods

### 1. Structures of Glycoprotein-Antibody

We start with available crystal structures for two Ebola (GPs) bound to their respective antibodies (ABs). One is Zaire strain GP-AB complex (PDB:3CSY)^11^, obtained from Zaire survivors and commonly called EBOV. In the trimeric structure of EBOV GP the GP1 and GP2 segments are bound to a neutralizing AB, KZ52, which is extracted from a human survivor of the 1995 Kikwit outbreak. The second structure is SUDAN GP-engineered antibody complex (16F6), (PDB: 3S88)^12^. 16F6 is also a neutralizing antibody bound to SUDAN GP. A noteworthy point is that major features of the tertiary backbone structures of both the GP and the neutralizing antibody are conserved between the Sudan and the Zaire strains, as seen in Figures 2c and 2d. However, the anti-Sudan AB is not effective against the Zaire strain, nor is the anti-Zaire AB effective against the Sudan strain. An important difference must be, therefore, at the level of side-chain interactions. We used these two structures as the template for our bioinformatics analysis and as a starting point for molecular dynamics (MD) simulations (Figure1a). For Zaire, we used only one of the GP-AB fragments (chains I, J, A, and B) (Figure1a), since the other 2 fragments are identical in the whole structure.

**Figure 1.**
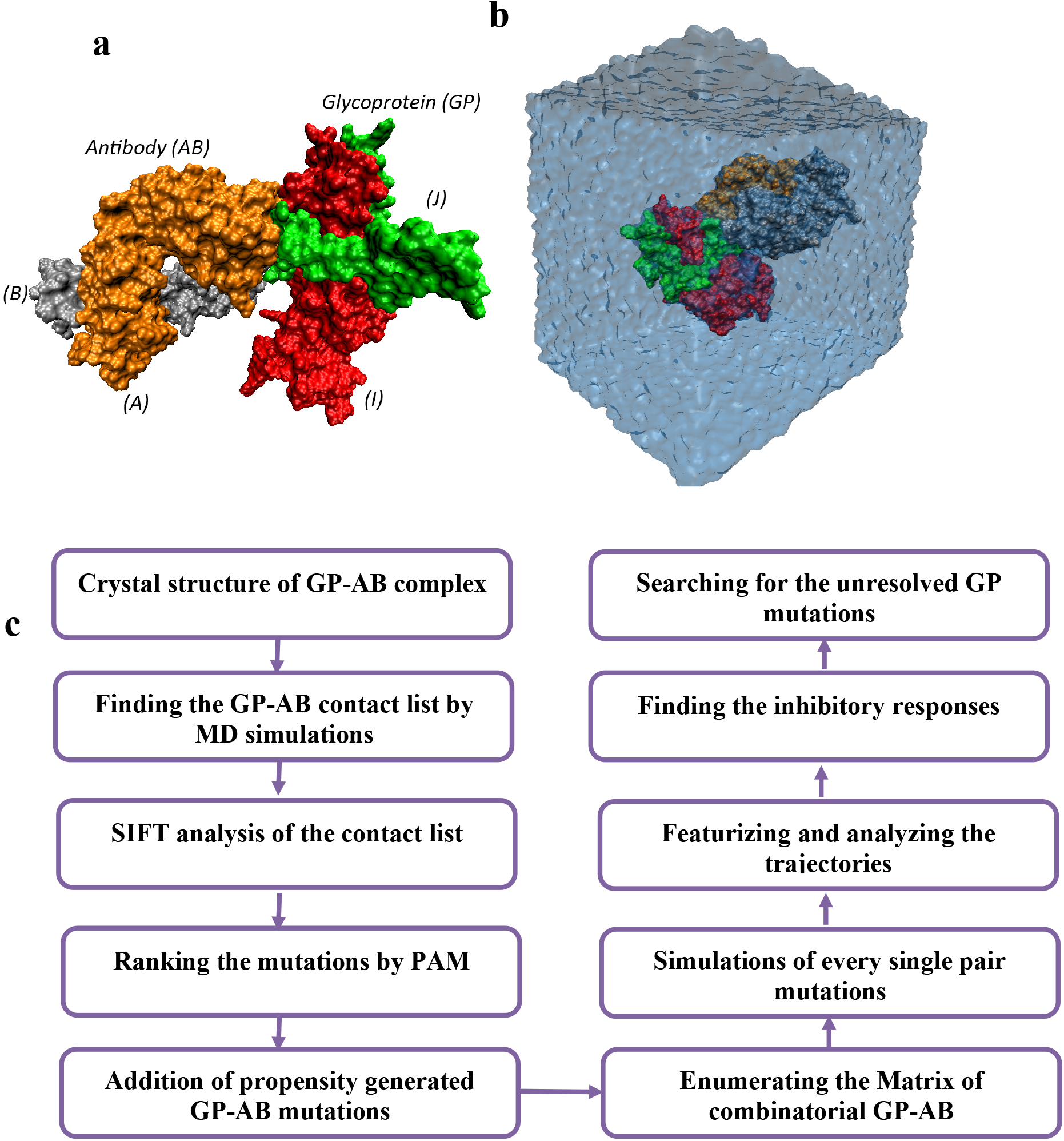
a) The one segment (out of chimeric of Glycoprotein (GP) -Antibody (AB) complex taken from Zaire Ebola virus crystal structure (PDB: 3CSY), we used chains I (in red),J (in green) (GP) and A (in orange), B (in grey) (AB) in our simulations. b) the solvated structure of GP-AB complex of Zaire (chains I,J,A,B of PDB:3CSY) used in our simulations. c) algorithm we used for reducing the mutagenesis study of GP-AB complex, featurization and generation of contact list and finding the inhibitory responses.

**Figure 2.**
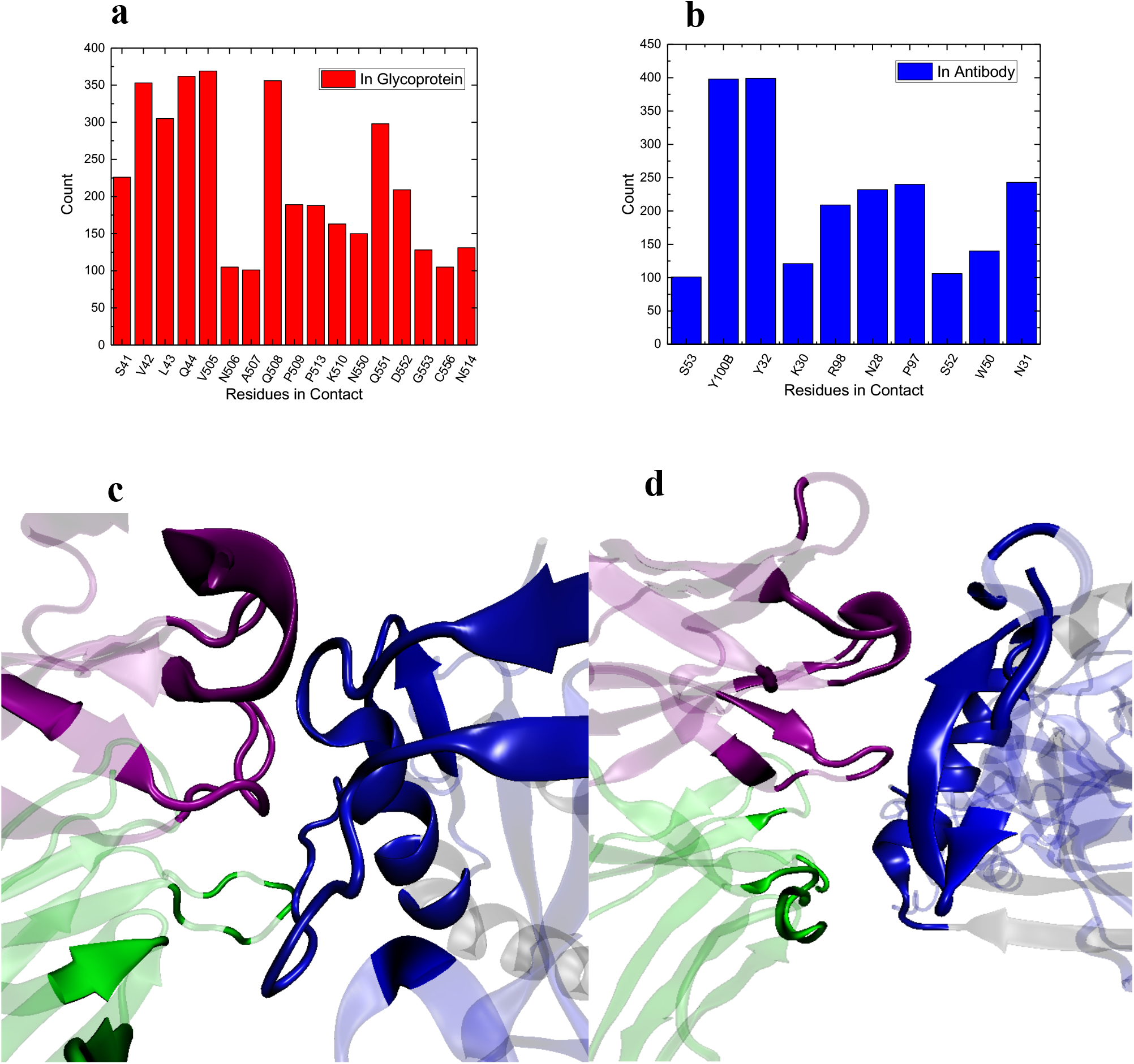
a) Important contact residues within 5 Å of antibody in glycoprotein (Zaire) computed by counting the residues in simulation trajectories. b) important contact residues within 5Å of glycoprotein in antibody (Zaire). The similar plots for SUDAN can be found in supporting information. c) KZ52 (antibody) and glycoprotein contact region. The contact residues within 5 Å were highlighted at the contact and the rest of the structure is shown as transparent. d) The contact residues within 5 Å were highlighted at the contact and the rest of the structure is shown as transparent for 16F6 contact residues with SUD GP.

### 2. Molecular Dynamics Simulations

We performed extensive MD simulations with NAMD 2.6 using the petascale Blue Waters machine.^13^ A typical simulation set up consisting of the protein, water and ions is shown in Figure 1b. The protein (GP-AB complex) is solvated in a water box extending 1.5 nm from the maximum and minimum of the protein, in each direction. We used the CHARMM27 force field^14^ parameters for all the proteins, and TIP3P water molecules. The SHAKE algorithm was used to maintain the rigidity of the water molecules. Periodic boundary condition was applied in all the three directions. The cutoff distance for the LJ interactions was 15 Å, and the long-range electrostatic interactions were computed using the Particle-Mesh-Ewald (PME) method.^15^ The time step was 1 fs. For each simulation, energy minimization was performed for 100,000 steps. Systems were then equilibrated for 3 ns with NPT ensemble at 1 atm pressure and 310 K temperature, ensuring that the simulated system was at physiological temperature and pressure. The system volume was free to change in the NPT ensemble but in fact did not change significantly during the simulation. Temperature was maintained at 310 K by applying the Langevin thermostat with a time constant of 0.1 ps.^16^ To equilibrate the WT structure of GP-AB, the two related wild type (WT) simulations of GP-AB for SUDAN and ZAIRE were run for 150 ns. Other mutant simulations were run for at least 12 ns after the equilibration. The trajectories were saved to the disk every 5 ps.

### 3. Contact List

To identify the most effective residues in the contact region of GP-AB complex, we performed long time-scale MD simulations on the complex structures depicted in Figure 1a, for both the Zaire and the Sudan GP-AB complexes in their wildtype forms. Instead of using a single snapshot reported in the crystal structure, we use MD trajectories for this analysis, which enabled us to identify the thermodynamically stable residues in the contact region of the complex (the contact list). The measure used to define the residues significant for interaction in both GP and AB was a 5-Å heavy atom distance between a residue in GP and a residue in AB, and vice versa. For the 120 ns of production trajectories (we discarded the initial 30 ns of equilibration simulation, out of 150 ns simulation mentioned in MD section), the interacting residues were counted 400 times (every 300 ps), and all residues which appeared at least in 100 counts were included in the contact list. The GP contact residues in Zaire are 17 (Figure 2a) and the AB contact residues in the same Zaire structure are 13 (Figure 2b). The frequency of the appearance of the residues in the contact list also indicates their level of importance in binding (Figure 2a, 2b). The proximities of the residues have also been discussed in the studies describing the crystal structures (Figure 2c, 2d).^11, 17^ We performed the same calculations for the Sudan structure to characterize the contact residues (For Sudan contact lists, see the Supporting Information)

### 4. Identifying the Most Probable Mutations

The pure combinatorics of varying the pairs of amino acids at the 17 GP and 13 AB sites makes it computationally prohibitive to test all mutations at random for favorable interactions. Thus, we needed to short-list the most probable mutations that the viral GP undertakes and the possible inhibitory mutations that the antibody would need to undertake to restore favorable association. The overall recipe for the computational design of antibodies is shown in Figure 1c, and described in detail below.

The first iteration of short-listing the mutations that the viral GP can undertake involves the Sorting Intolerant From Tolerant (SIFT) algorithm, which uses large databases of mutations and their functional consequences to identify substitution mutations that are most likely to preserve function.^18^ SIFT analysis was performed on all fragments of GP and AB in both Zaire and SUDAN structures (Supporting Information SIFT.xslc). Combining the contact lists (see the Contact List section above) obtained from MD simulations for the GP-AB interface and the SIFT results, we obtained the shortlisted mutations that both GP (Table 1) and AB (Table 2) can plausibly perform. In order to rank and prioritize the significant mutations both in GP and AB, we used the Position Acceptable Mutations (PAM) to identify which of the mutations that were obtained from SIFT (see Supporting Information, PAM.xslx) are most likely.^8^ PAM ranks the most probable mutations based on the natural frequency of their occurrence in sequence and structure databases.^8^ We categorized the ranking into three sections: highly probable (red color), probable (orange) and medium probability (blue) based on the PAM scores (See Table 1 and Table 2 for GP and AB mutations and the colors denoting the order of importance). In order to not miss the highly interacting residues at the contacts irrespective of their ranking with PAM, we also included the highly interacting residues based on propensity for occurring specifically at antibody-antigen interfaces^19^ (see propensity.xslx in Supporting Information). These mutations are designated in grey color in Tables 1 and 2. Now that we have the possible mutations and ranked them based on their significance for both GP and AB contact residues, we can generate the table of possible simulations. Before doing this, we need to know what residues in the GP contact list interacts with which residues in the AB contact list. To identify this, we looked at the hydrogen bonds, salt bridges and the initial interaction matrix in the crystal structure of GP-AB complex. The interaction matrix we built is consistent with the crystal structure interaction matrix shown in Ref.^11^ For clarity, we show the interaction matrix in Table 3 for Zaire. In this table, the first two columns list the GP and AB contact residues, respectively. The residues in each row interact with each other.

**Table 1:**
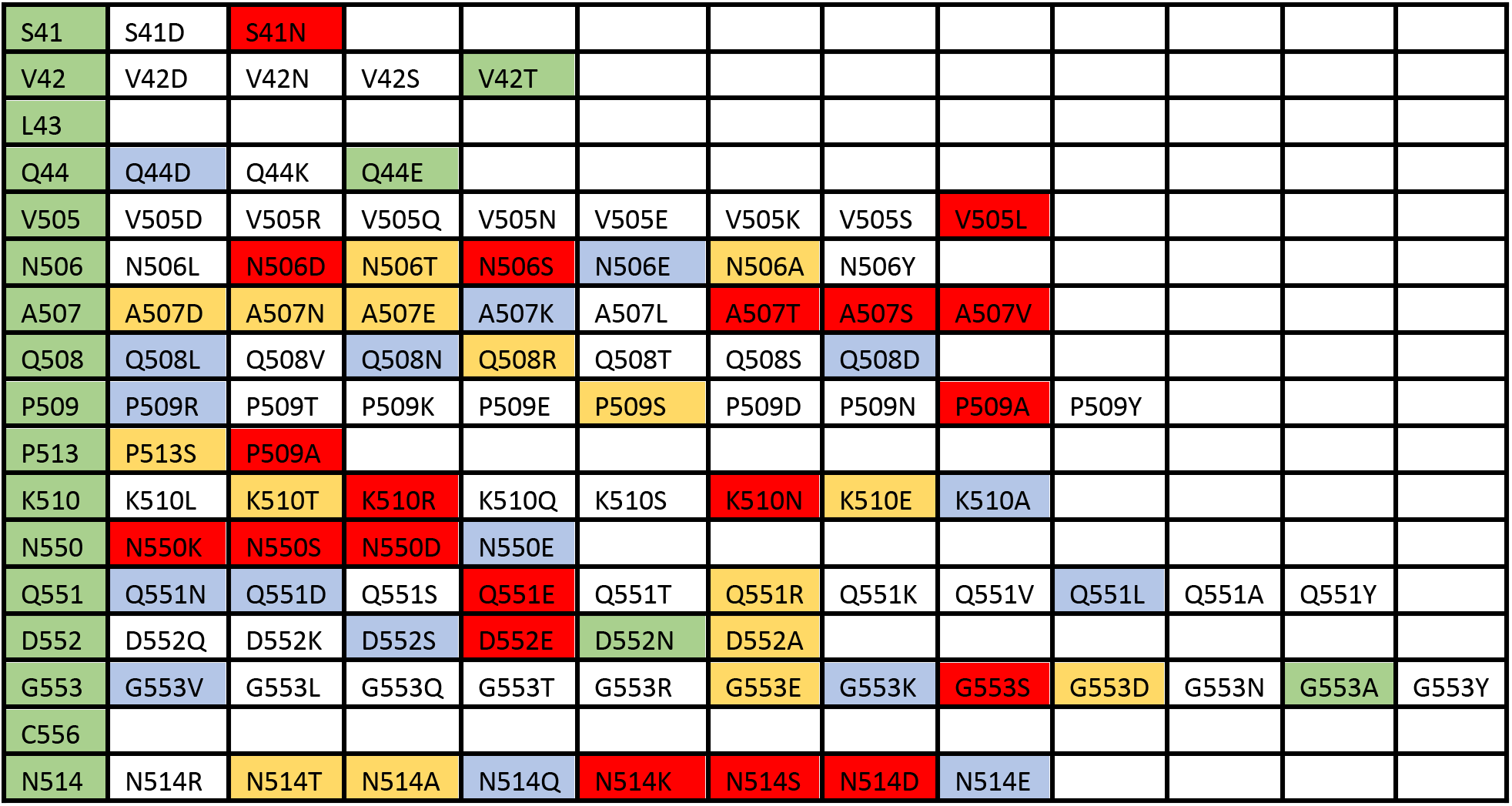
The tolerable mutations that each amino acid of GP contact can perform for Zaire strain. The first column shows the GP contact amino acids.

|  |  |  |  |  |  |  |  |  |  |  |  |  |
| --- | --- | --- | --- | --- | --- | --- | --- | --- | --- | --- | --- | --- |
| S41 | S41D | S41N |  |  |  |  |  |  |  |  |  |  |
| V42 | V42D | V42N | V42S | V42T |  |  |  |  |  |  |  |  |
| L43 |  |  |  |  |  |  |  |  |  |  |  |  |
| Q44 | Q44D | Q44K | Q44E |  |  |  |  |  |  |  |  |  |
| V505 | V505D | V505R | V505Q | V505N | V505E | V505K | V505S | V505L |  |  |  |  |
| N506 | N506L | N506D | N506T | N506S | N506E | N506A | N506Y |  |  |  |  |  |
| A507 | A507D | A507N | A507E | A507K | A507L | A507T | A507S | A507V |  |  |  |  |
| Q508 | Q508L | Q508V | Q508N | Q508R | Q508T | Q508S | Q508D |  |  |  |  |  |
| P509 | P509R | P509T | P509K | P509E | P509S | P509D | P509N | P509A | P509Y |  |  |  |
| P513 | P513S | P509A |  |  |  |  |  |  |  |  |  |  |
| K510 | K510L | K510T | K510R | K510Q | K510S | K510N | K510E | K510A |  |  |  |  |
| N550 | N550K | N550S | N550D | N550E |  |  |  |  |  |  |  |  |
| Q551 | Q551N | Q551D | Q551S | Q551E | Q551T | Q551R | Q551K | Q551V | Q551L | Q551A | Q551Y |  |
| D552 | D552Q | D552K | D552S | D552E | D552N | D552A |  |  |  |  |  |  |
| G553 | G553V | G553L | G553Q | G553T | G553R | G553E | G553K | G553S | G553D | G553N | G553A | G553Y |
| C556 |  |  |  |  |  |  |  |  |  |  |  |  |
| N514 | N514R | N514T | N514A | N514Q | N514K | N514S | N514D | N514E |  |  |  |  |

**Table 2:** The tolerable mutations that each amino acid of AB contact can perform for Zaire strain. The first column shows the AB contact amino acids.

|  |  |  |  |  |  |  |  |  |  |  |  |  |
| --- | --- | --- | --- | --- | --- | --- | --- | --- | --- | --- | --- | --- |
| S53 | S53P | S53D | S53Q | S53R | S53N | S53L | S53Y |  |  |  |  |  |
| Y100B | Y100D | Y100N | Y100L | Y100Q | Y100V |  |  |  |  |  |  |  |
| Y100B | Y100D | Y100N | Y100L | Y100Q | Y100V |  |  |  |  |  |  |  |
| R98 | R98P | R98D | R98K | R98Q | R98N | R98E | R98T | R98S | R98L | R98Y |  |  |
| Y32 | Y32Q | Y32K | Y32E | Y32R | Y32D | Y32L | Y32N | Y32T | Y32S |  |  |  |
| R98 | R98P | R98D | R98K | R98Q | R98N | R98E | R98T | R98S | R98L | R98Y |  |  |
| K30 | K30E | K30T | K30Q | K30S | K30R |  |  |  |  |  |  |  |
| R98,W50 | R98P | R98D | R98K | R98Q | R98N | R98E | R98T | R98S | R98L | R98Y |  |  |
| R98 | R98P | R98D | R98K | R98Q | R98N | R98E | R98T | R98S | R98L | R98Y |  |  |
| N28 | N28K | N28R | N28T | N28S |  |  |  |  |  |  |  |  |
| W50 | W50P | W50D | W50Q | W50N | W50K | W50E | W50R | W50R | W50T | W50S | W50L | W50Y |
| P97 | P97D | P97Q | P97K | P97R | P97E | P97T | P97N | P97S | P97L | P97Y |  |  |
| S53 | S53P | S53D | S53Q | S53R | S53N | S53L | S53Y |  |  |  |  |  |
| S52 | S52P | S52D | S52E | S52K | S52N | S52Q | S52T | S52R | S52L | S52Y |  |  |
| W50 | W50P | W50D | W50Q | W50N | W50K | W50E | W50R | W50R | W50T | W50S | W50L | W50Y |
| P97 | P97D | P97Q | P97K | P97R | P97E | P97T | P97N | P97S | P97L | P97Y |  |  |
| N31 | N31Q | N31K | N31E | N31R | N31T | N31S | N31D |  |  |  |  |  |
| W50 (2nd) | W50P | W50D | W50Q | W50N | W50K | W50E | W50R | W50R | W50T | W50S | W50L | W50Y |

**Table 3:** The interaction matrix between Amino acids in GP and AB, in Zaire structure. For example, Q508 in GP interacts with both R98 and W50.

| GP | AB |
| --- | --- |
| S41 | S53 |
| V42 | Y100B |
| L43 | Y100B |
| Q44 | R98 |
| V505 | Y32 |
| N506 | R98 |
| A507 | K30 |
| Q508 | R98,W50 |
| P509 | R98 |
| P513 | N28 |
| K510 | W50 |
| N550 | P97 |
| Q551 | S53 |
| D552 | S53,S52 |
| G553 | W50 |
| C556 | P97 |
| N514 | N31 |

Based on Tables 1, 2 and 3 (GP mutations, AB mutations and the interacting residues), different combinational mutated GP-AB complexes emerge as suitable for detailed analysis by MD simulation. The populated simulations are those where the corresponding cells in both Tables 1 and 2 have a non-zero value. 1,270 simulations were thus set up and run for mutated GP-AB complexes, 734 for Zaire and 536 for Sudan. Altogether, 13.4 μs of simulation time was performed.

## Results and Discussions

To test the viability of our method and the appropriateness of feature selection on the MD trajectories analysis, we first did MD simulations of four combinations of viral protein associated with antibody; namely, Zaire GP to Zaire AB (GP(Z)-AB(Z)), Sudan GP to Sudan AB (GP(S)-AB(S)), Zaire GP to Sudan AB (GP(Z)-AB(S)), and Sudan GP to Zaire AB (GP(S)-AB(Z)). As a prelude to the latter two simulations, we structurally aligned the AB of Sudan to AB of Zaire (creating GP(Z)-AB(S) simulation set) and also GP of Zaire to GP of Sudan (creating GP(S)-AB(Z)) using VMD.^20^ For GP(Z)-AB(Z) we used the same contact list tabulated in Table 3. For GP(S)-AB(Z), GP(Z)-AB(S) and GP(S)-AB(S)simulations, we generated the contact list as described in Section 3. We ran each simulation for 150 ns. We computed the root mean squared displacement (RMSD) of all heavy atoms in residues in contact list and the contact distances of the heavy atoms of residues in contact list for each of the four cases (Figure 3a). The average contact distances and RMSD (noted in the same order in bracket, all in Å) are: GP(Z)-AB(Z): [2.44, 4.42], GP(S)-AB(S): [2.34, 4.01], GP(Z)-AB(S): [6.47, 7.78], GP(S)-AB(Z): [5.87, 7.45]. Very low RMSD, contact distances and their low standard deviations in the wildtype structures depict the stability and strong binding at the GP and AB at their interface. However, the GP(S)-AB(Z) and GP(Z)-AB(S) binding are weak, as shown by the high average contact distance and RMSD (Figure 3). Next, we computed the interaction energies of the interface. We computed the interaction energies of pairs in the contact list for each frame and averaged it over the production run. The computed interaction energies for the contact residues GP(Z)-AB(Z): [12.36 kcal/mol], GP(S)-AB(S): [12.25 kcal/mol], GP(Z)-AB(S): [2.78 kcal/mol], GP(S)-AB(Z):[3.11 kcal/mol] show the stability of the native complexes versus the swapped ones, i.e., GP(Z)-AB(S) and GP(S)-AB(Z).

**Figure 3.**
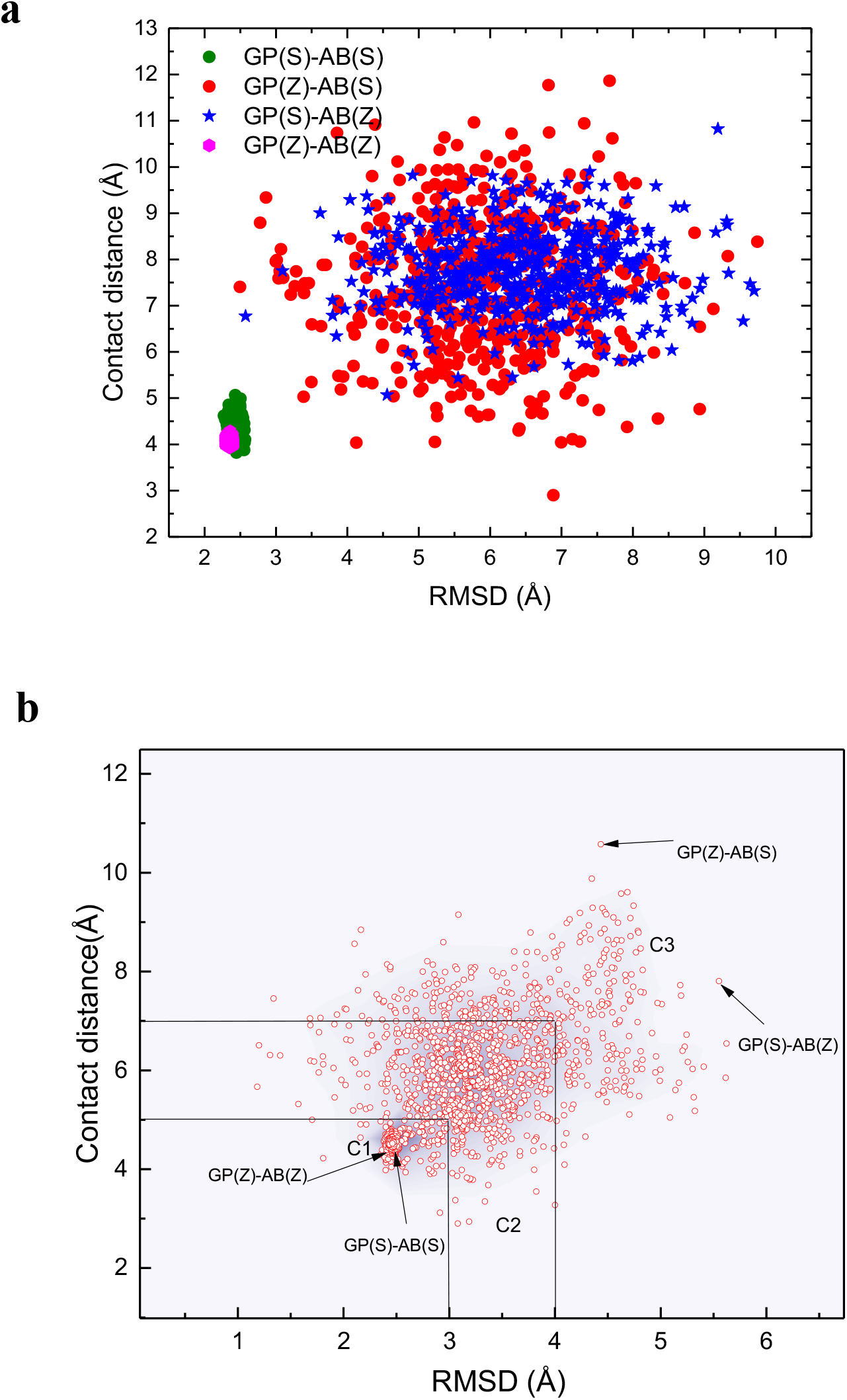
a) The benchmark of featurization based on RMSD and contact residue distances for GP(Z)-AB(Z), GP(S)-AB(S), GP(Z)-AB(S), GP(S)-AB(Z). b) RMSD and contact distances of 1273 mutations for Zaire and Sudan (Note the relative location of wildtype structures with mutations). 1270 simulations are point mutations and the additional three are (GP(S)-AB(S), GP(Z)-AB(S), GP(S)-AB(Z). Three levels of responses, C1:strong, C2: mild and C3: weak interactions are shown. The lower RMSD and contact distance represent higher and stronger interaction and binding

We mapped the RMSD and contact distances of 1,273 GP-AB pair mutations of both Sudan and Zaire complexes and categorized the RMSD and contact distances into three groups (Figure 3b). These three groups are based on these selection criteria (C1: RMSD < 3 Å & contact distance < 5 Å; C2: 3 <RMSD <4 & 5 < contact distance <7, C3: RMSD > 4 & contact distance > 7). C1 pair mutations are very stable and likely would offer high protection against infection, C2 mutations are moderately stable, and C3 are definitely unstable and would likely not inhibit infection.

To better understand the binding energetics at the GP-AB interface, we computed the interaction energy between the residues in the contact list (Table 3). In case of a mutation, the mutated residues in Table 3 were updated in the list and calculations were performed on the updated contact list. The interaction energy, contact distance and RMSD of Zaire are shown in a 3D plot in Figure 4 with the projections of pair plots on each plane. As the RMSD and contact distance decrease, the magnitude of the interaction energy increases, which represents stronger binding. In general, there is an inverse relation between RMSD and contact distances on the one hand, and interaction energy on the other hand. The corresponding searchable data is available in Supporting Information (Zaire_all_mutations.xlsc). A total of 75 viral mutations were tested. Of the 75, our computational procedure succeeded in producing a mutated antibody with a more favorable interaction energy than 12.0 kcal/mol for 64 of them. This criterion for “certain success” was chosen because we computed the interaction energy for the native WT-WT to be 12.36 kcal/mol, and that antibody is known to be effective in vivo. The 64 mutation pairs with predicted successful inhibitory responses are tabulated in Table S1 in supporting information. It is seen that all RMSD values in this table are 3.03 Å or less. We found certain mutations in AB that are predicted to make the current GP(Z)-AB(Z) binding even stronger than Zaire WT-WT structure. These mutations are WT-S53L and WT-R98D with interaction energies of 12.82 and 13.89 kcal/mol, respectively. Both energy values are greater in magnitude than WT-WT interaction energy (12.36 kcal/mol). For the 11 viral strains that our procedure for mutating antibody did not meet the 12 kcal/mol criterion, the weakest affinity was 10.5 kcal/mol At this writing we are not certain whether the constraints of our search algorithm for antibody mutations, instituted for computational efficiency and shown in Table 2, prevented us from finding a more effective binding antibody in these 11 instances, nor whether 10.5 kcal/mol might be enough to be effective in vivo.

**Figure 4.**
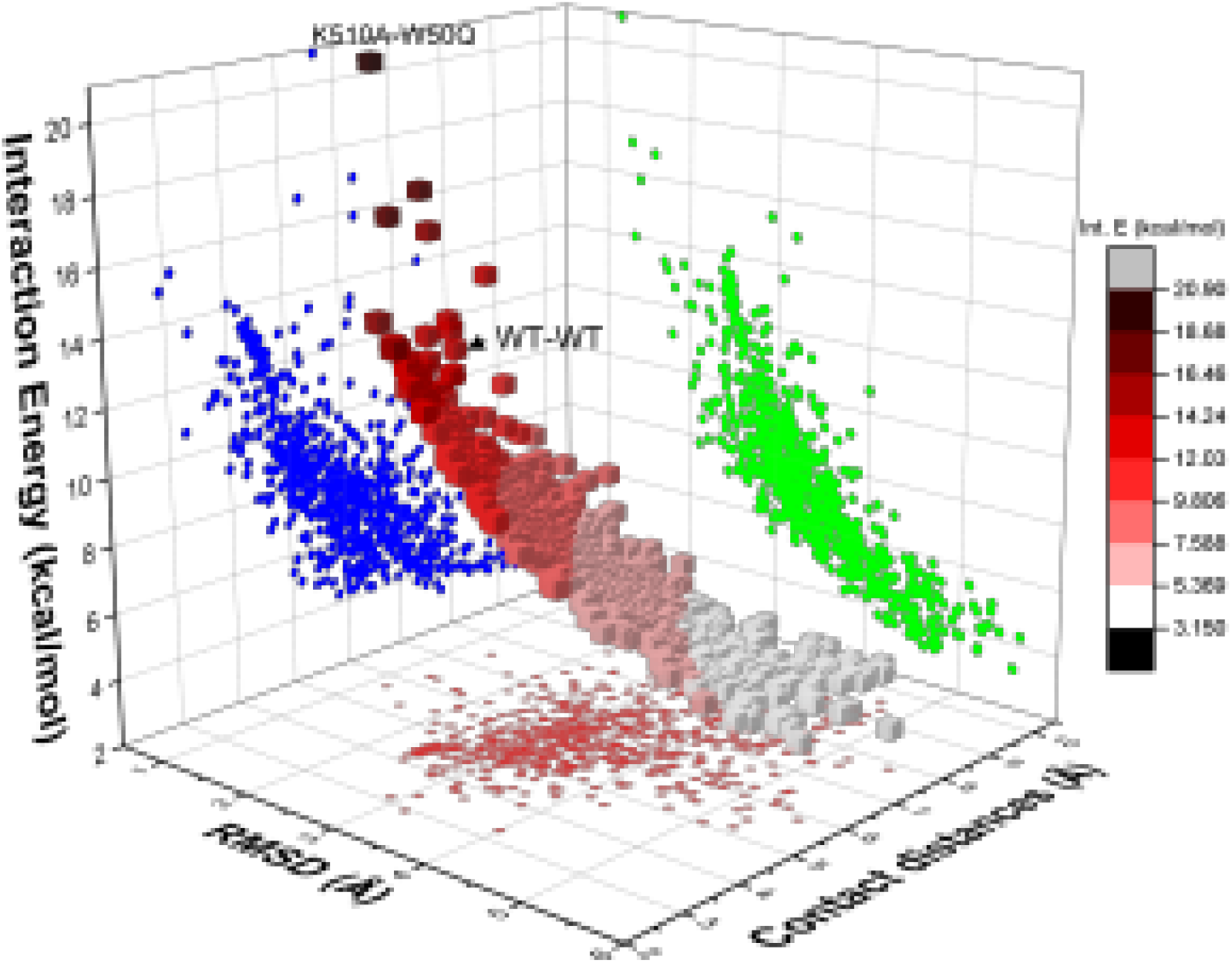
The 3D plot of RMSD, contact distance and interaction energy of all Zaire mutations. The WT-WT and the highest interaction energy pairs (K510A-W50Q) are marked. The darker colors represent higher interaction energies, lower RMSD and contact distances in the 3D markers. The projection plots of interaction energy-contact distances (in blue color), RMSD-contact distances (in red), and RMSD-interaction energy (in green) are shown in three orthogonal planes.

For all of the combinations of GP mutations and varying AB mutations we computed the RMSD, contact distances and interaction energies for each. Examples are shown graphically in Figure 5. For example, Figure 5d shows that for V505N, the trial AB mutations selected as described in the Methods section were Y32L, Y32K, Y32E, Y32Q, Y32R and Y32D. The order of response based on the magnitude of the interaction energies are Y32D>Y32R>Y32Q>Y32K>Y32E>WT>Y32L (Figure 5d). In this case, Y32D, the most energetically favorable of the AB response mutations, also shows the minimum contact distance and RMSD. The best response for V505N is Y32D, while the KZ52 wild type antibody has a very weak binding interaction (Figure 5d). For those GP mutation, a responding AB mutation with more favorable interaction energies than 10.5 kcal/mol has been found. For 11 GP mutations out of 75 mutations, our automated method did not find an interaction energy reaching the 12.0 kcal/mol cutoff. In all of those cases an energy of at least 10.5 kcal was achieved. A judgment of whether 10.5 kcal/mol would be adequate to provide immunity in the biological context is outside the scope of this study; it may be that 10.5 kcal/mol is good enough. It may also be that our automated criteria for which AB mutations are tolerable or preferred was too stringent, and that a more exhaustive search would provide more favorable association energies.

**Figure 5.**
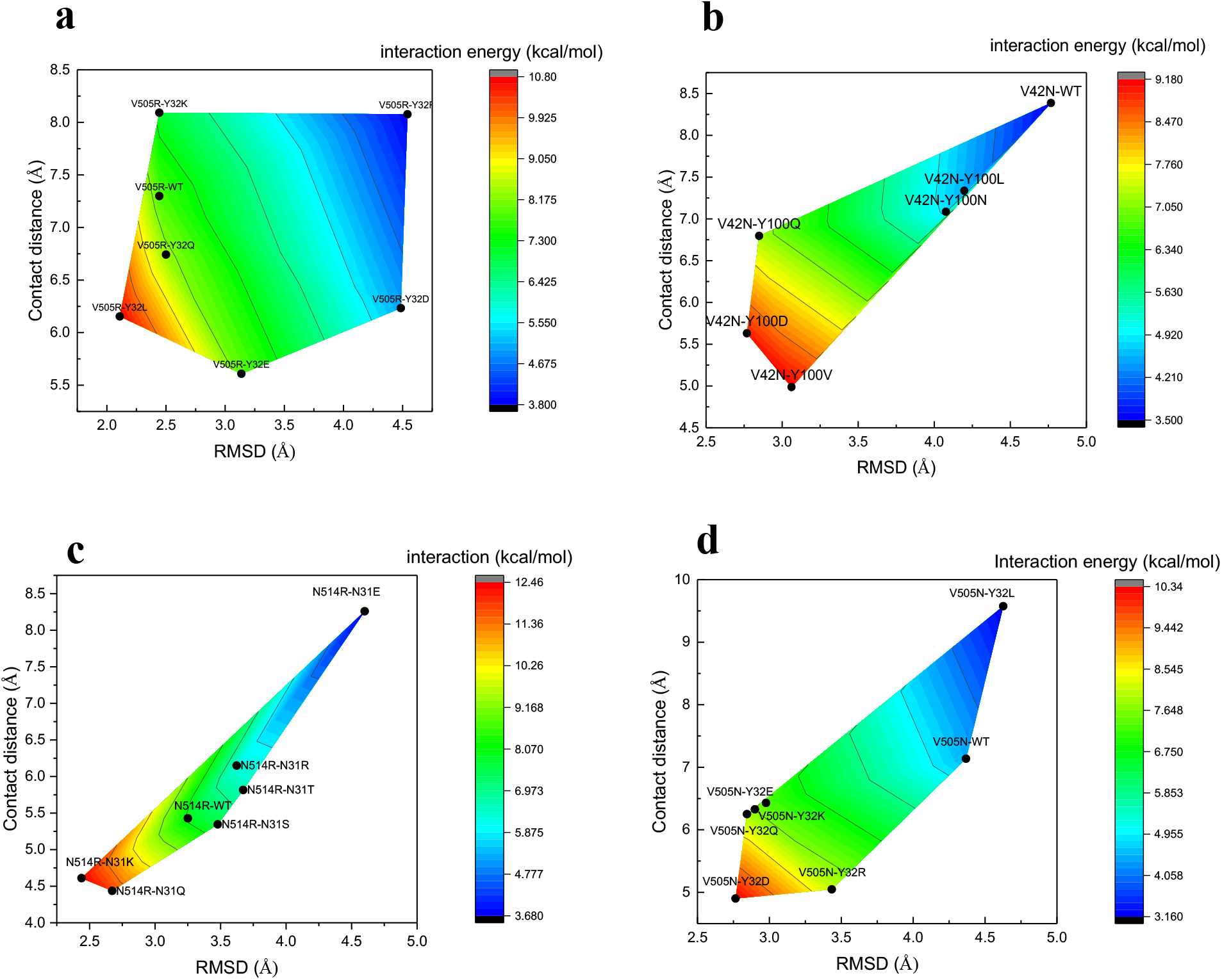
The AB mutation responses to GP mutations and presentation of the contact distances, RMSD and interaction energies. The best response has the lowest RMSD and contact distance and highest binding energy. These four selections are based on the wide range of AB responses to their respective mutations. a) V505R, b) V42N, c) N514R, d) V505N.

The results of an alanine-scanning study on Ebola Zaire GP match very well with our prediction of viral escape mutations.^21^ Davidson et al. mutated each residue of the Zaire GP to alanine and measured changes in GP-KZ52 binding affinities relative to the WT. They found C511, N550, D552, G553, and C556 to be critical in binding to KZ52, which is consistent with our findings of the important contact amino acids. The viral escape mutations are available in SI (Escape_Zaire_mutations.csv).

### Multiple mutation effect and antibody response

Since viruses like Ebola would sometimes not escape the antibodies with single mutation in GP, and since mutations occur at more than one position of the viral protein, it is necessary to also study the effect of multiple mutations which may simultaneously occur in the GP. As a first step toward this goal, we set up simulations with 2 mutations on GP (N514R, V42N, in Zaire strain) occurring simultaneously. The single point mutation data shows that V42N-Y100V, N514R-N31Q produce energetically stable responses (Figure 5). The first simulation that we ran includes both N514R-WT and V42N-WT, which resulted in an interaction energy of 3.12 kcal/mol, implying an unstable interface. Note that the interaction energies for single mutant simulations for N514R-WT and V42N-WT were 7.93 and 3.50 kcal/mol, respectively. For the second simulation, we tested the N514R-N31Q and V42N-Y100V combination. The computed interaction energy for this second simulation is 10.13 kcal/mol which is between the value for single mutation simulation of N514R-N31Q and V42N–Y100V (Figure 5b and Figure 5c). The N31Q and Y100V mutants have the most favorable interaction energies with N514R and V42N GP mutations, respectively. These examples support the conclusion that the restoring of multiple simultaneous mutation is possible by iterating the procedure for a single mutation.

## Conclusion

We have developed a computational method for high throughput screening of synthetic antibodies to inhibit the next generation of viral epidemics. In real life this method would be based on sequencing the emergent virus. In our work we use statistics of amino acid substitutions to project the likely sequences of new viral strain GPs. We believe the combination of bioinformatics, structural biology, and simulations can help us to screen and analyze the vast population of possible mutations on both viral proteins and possible candidates for synthetic antibodies. For Ebola virus, we projected the most likely viral mutations which disrupt the binding interface with the current neutralizing antibody and the possible synthetic antibodies that restore binding. Our approach can be widely applied to other viruses where structures of viral coat protein-antibody pairs can be obtained. We envisage in the future the following pathway to proactively create synthetic antibodies against variants of the virus: 1) obtain a structure for the viral coat protein bound to an antibody, 2) use the method described in this paper to project likely mutations in the viral coat protein in future strains, and to compute the sequences in the binding region of the antibody that will successfully counter the likely mutations, and 3) Using available methods for engineering antibodies, create synthetic antibodies with the desired sequence in the antibody binding region. We look forward to, and would be happy to collaborate with, experimental projects to test our methods.

