## Supporting Information for "Computational Recipe for Designing Antibodies against the Ebola Virus"

| Simulation ID | MUTATIONS | RMSD | Contacts | Interaction E |
|---|---|---|---|---|
| 5 | WT-S53L | 2.42161 | 4.50906 | 12.82146998 |
| 50 | WT-R98D | 2.40356 | 4.19118 | 13.89750231 |
| 51 | WT-R98K | 2.41192 | 4.69624 | 12.35989706 |
| 60 | Q44D-R98N | 1.7216 | 6.17876 | 13.16116868 |
| 62 | Q44K-WT | 2.3991 | 4.45682 | 13.09346497 |
| 86 | V505D-Y32R | 2.46923 | 4.55969 | 12.43458151 |
| 87 | V505D-Y32D | 2.43419 | 4.61798 | 12.45436284 |
| 123 | V505K-Y32L | 2.36577 | 4.38769 | 13.48713111 |
| 143 | N506L-R98N | 3.1881 | 2.93901 | 14.94152881 |
| 173 | N506A-WT | 2.49261 | 4.2943 | 13.07920424 |
| 181 | N506Y-R98P | 1.687 | 6.19233 | 13.40166817 |
| 185 | N506Y-R98N | 2.43161 | 4.34457 | 13.25217969 |
| 231 | A507S-K30Q | 1.4013 | 6.30239 | 15.85227652 |
| 233 | A507S-K30R | 2.85408 | 3.91738 | 12.52178353 |
| 245 | Q508L-R98N | 2.43146 | 4.63199 | 12.43063439 |
| 252 | Q508V-R98N | 2.49943 | 4.4043 | 12.7177465 |
| 258 | Q508N-R98Q | 2.50567 | 4.46999 | 12.49964304 |
| 287 | Q508D-R98N | 2.47678 | 4.59949 | 12.28940692 |
| 295 | P509R-R98E | 2.4536 | 4.40959 | 12.93975524 |
| 304 | P509K-R98P | 1.97988 | 5.80649 | 12.1779864 |
| 339 | P509A-R98P | 2.45113 | 4.37515 | 13.05475552 |
| 340 | P509A-R98D | 2.43187 | 4.30238 | 13.38070248 |
| 356 | P513S-WT | 2.41608 | 4.66131 | 12.43107649 |
| 360 | P513S-N28S | 2.54704 | 4.5679 | 12.03304892 |
| 362 | P509A-N28R | 2.72905 | 4.16347 | 12.3214297 |
| 377 | K510L-W50E | 2.09021 | 5.18886 | 12.90821413 |
| 379 | K510T-W50P | 2.45648 | 4.33243 | 13.15476968 |
| 395 | K510Q-W50Q | 2.45537 | 4.57566 | 12.46112763 |
| 423 | K510A-W50Q | 1.1835 | 5.66688 | 20.8744844 |
| 425 | K510A-W50K | 3.05763 | 3.37032 | 13.58538552 |
| 434 | N550K-P97D | 2.45967 | 4.60491 | 12.3603294 |
| 440 | N550S-WT | 2.42622 | 4.43542 | 13.00957493 |
| 465 | Q551N-S53R | 2.46399 | 4.39711 | 12.9217628 |
| 467 | Q551N-S53L | 2.4441 | 4.40268 | 13.01043879 |
| 485 | Q551E-S53Q | 2.52357 | 4.43227 | 12.51660282 |
| 495 | Q551T-S53L | 2.43088 | 4.528 | 12.71915023 |
| 497 | Q551R-S53P | 2.40625 | 4.33427 | 13.42367185 |
| 512 | Q551V-S53D | 2.80693 | 4.00103 | 12.4659289 |
| 516 | Q551V-S53L | 2.44286 | 4.58563 | 12.49771013 |
| 545 | D552Q-S52P | 2.21566 | 4.90369 | 12.88551886 |
| 546 | D552Q-S52D | 2.48531 | 4.66908 | 12.06468961 |
| 548 | D552Q-S52K | 2.40397 | 4.48363 | 12.98880586 |

| | | | | |
|---|---|---|---|---|
| 559 | D552S-S52P | 2.45335 | 4.04207 | 14.11772427 |
| 580 | D552A-S52P | 2.38261 | 4.56686 | 12.86640973 |
| 591 | G553V-W50K | 2.83344 | 3.95834 | 12.48248118 |
| 606 | G553Q-W50E | 1.80854 | 6.12242 | 12.64377614 |
| 617 | G553R-W50Q | 1.57306 | 5.31885 | 16.73266072 |
| 622 | G553E-W50P | 2.01987 | 5.70165 | 12.15637426 |
| 626 | G553E-W50K | 2.46514 | 4.25001 | 13.36277005 |
| 631 | G553K-W50Q | 2.43496 | 4.50169 | 12.77205028 |
| 635 | G553S-WT | 2.46455 | 4.43712 | 12.8023362 |
| 641 | G553S-W50E | 1.33882 | 7.45484 | 14.02708787 |
| 659 | G553A-W50Q | 2.9128 | 3.11671 | 15.42129963 |
| 665 | G553Y-W50D | 2.39962 | 4.51192 | 12.9307636 |
| 668 | G553Y-W50K | 2.48643 | 4.3 | 13.09433185 |
| 671 | WT-N31K | 2.2138 | 5.21743 | 12.12084846 |
| 678 | N514R-N31K | 2.43919 | 4.60978 | 12.45094187 |
| 686 | N514T-N31E | 2.45864 | 4.3616 | 13.0553121 |
| 691 | N514A-N31Q | 2.43072 | 4.61202 | 12.48825964 |
| 697 | N514Q-WT | 1.91544 | 5.44999 | 13.41108076 |
| 707 | N514K-N31E | 1.30524 | 6.30922 | 17.00051156 |
| 717 | N514S-N31S | 2.49853 | 4.54882 | 12.31812808 |
| 731 | N514E-N31S | 2.46718 | 4.32877 | 13.10879249 |

**Table S1**: The "certainly successful" inhibitory mutation responses for Zaire. The selection criteria is interaction energy> 12.0 kcal/mol, selected to be very close to WT(GP)-WT(AB) interaction energy of 12.34 kcal/mol.

**Contact residues in SUDAN GP-AB complex**

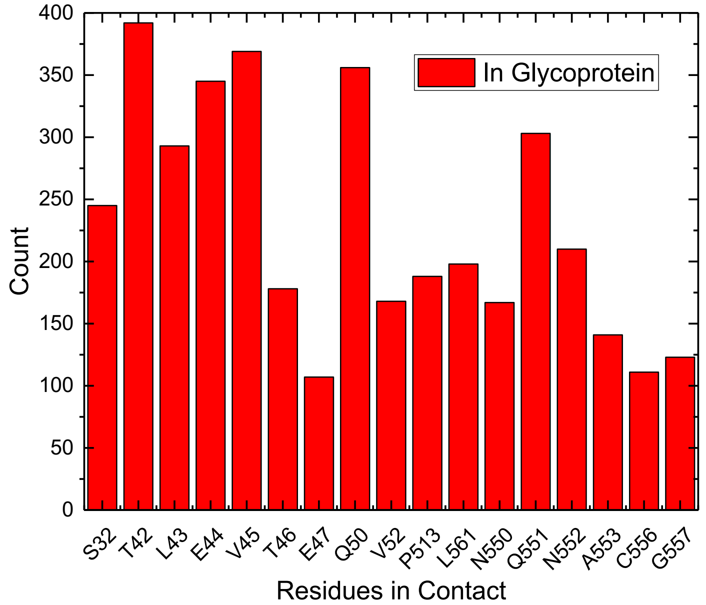


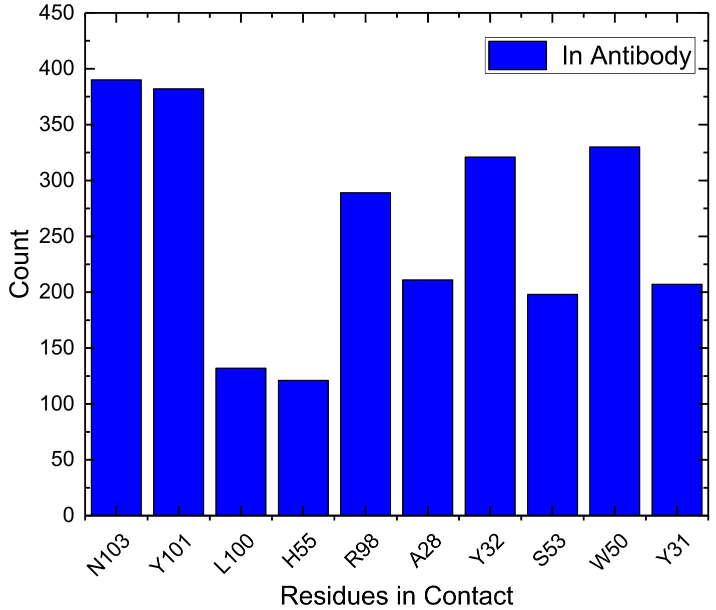


**Figure S.1**. a) Important contact residues within 5 Å of antibody in glycoprotein (SUDAN) computed by counting the residues in simulation trajectories. b) important contact residues within 5Å of glycoprotein in antibody (SUDAN).

**a**

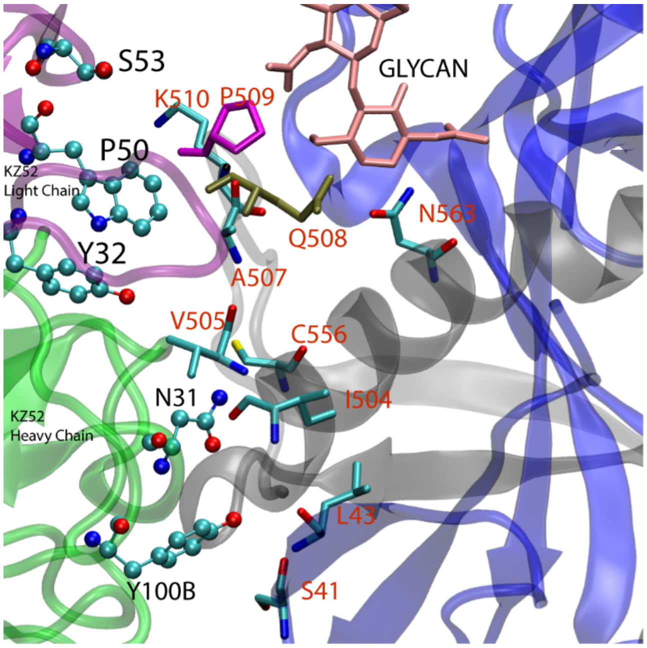


**b**

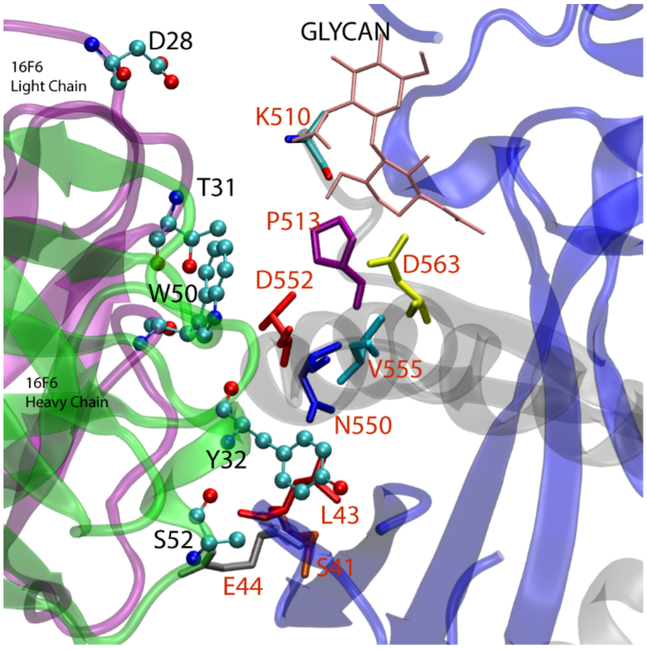


**Figure S2** a: KZ52 (antibody) and glycoprotein contact region and the key residues in binding site. b| 16F6 contact residues with SUD GP. The residues in AB are illustrated in CPK, and residues in GP are illustrated in Licorice representations in c and d subplots.
